# Hepatic stearoyl-CoA desaturase deficiency ameliorates hyperglycemia through bile acid signaling in an insulin-independent manner

**DOI:** 10.64898/2026.07.07.737046

**Authors:** Mugagga Kalyesubula, Daehan Kim, Woo Sung Kim, Nicole B. Wicker, Jonathan Williams, Veronica P. Christofi, Ethan Anderson, Jacqueline R. Miller, Dylan Cootway, Kaela Groppel, Daniel Bergman, Snehal N. Chaudhari, James M. Ntambi

## Abstract

Hyperglycemia in Type 1 Diabetes (T1D) is managed almost exclusively via exogenous insulin therapy, an approach restricted by significant glycemic fluctuations, long-term side effects such as weight gain, and high economic burden. Identifying physiological pathways capable of clearing blood glucose independent of insulin is therefore of paramount clinical importance. Here, we demonstrate that liver-specific stearoyl-CoA desaturase-1 (SCD1) deficiency protects against diabetic hyperglycemia and hepatic steatosis in an insulin-independent manner. SCD1 ablation decreases cellular oleate availability, altering lipid flux and redirecting excess cholesterol into alternative biosynthetic pathways. This redirection drives a 2-fold elevation in hepatic bile acids and a striking 10-fold increase in plasma bile acids, predominantly characterized by the accumulation of taurocholic acid. This shifted bile acid pool stimulates the expression of glucose transporter 1 (*Glut1*) in the liver via activation of the nuclear hormone receptor FXR, facilitating basal glucose clearance in the absence of insulin. Genetic deletion models show that while the hepatokine FGF21 serves as a partial mediator of this phenotype, the local bile acid-FXR axis remains a sufficient driver of systemic glucose clearance. Finally, we show that dietary oleate supplementation completely reverses this protective phenotype, turning down *Glut1* expression and restoring overt diabetes. Together, our findings uncover a novel bile acid-FXR-Glut1 signaling axis triggered by SCD1 inhibition, offering a framework for insulin-independent glycemic control.

**Article Highlights:** Here, we demonstrate that liver-specific stearoyl-CoA desaturase-1 (Scd1) deficiency protects against diabetic hyperglycemia and hepatic steatosis in an insulin-independent manner. Mechanistically, Scd1 ablation redirects excess cholesterol into bile synthesis, predominantly characterized by an increase in liver and plasma taurocholic acid. These shifted bile acids stimulate hepatic glucose transporter 1 (*Glut1*) expression via the farnesoid X receptor (FXR) activation to facilitate basal glucose clearance. While the hepatokine FGF21 acts as a partial systemic mediator, the local bile acid-Fxr axis remains a sufficient driver of clearance, a protective phenotype completely reversed by dietary oleate supplementation.

## Introduction

Hyperglycemia is the primary driver in the onset and progression of both Type 1 Diabetes (T1D) and Type 2 Diabetes (T2D). Prolonged elevation of blood glucose predisposes patients to devastating microvascular complications, such as retinopathy, nephropathy, and neuropathy, as well as macrovascular diseases (1–5). These cardiovascular complications represent the leading cause of morbidity and mortality among the diabetic population (2,6). While hyperglycemia in T2D stems from peripheral insulin resistance or insufficient pancreatic insulin secretion (7,8), T1D hyperglycemia is caused by absolute insulin deficiency following the autoimmune or idiopathic destruction of pancreatic β cells (9,10). Consequently, there is an urgent need for effective glycemic management strategies to mitigate these long-term cardiovascular risks.

Currently, T1D management relies heavily on exogenous insulin therapy, typically administered via multiple daily subcutaneous injections. However, insulin monotherapy has inherent limitations in achieving tight, physiological glycemic control. Extreme glucose fluctuations often lead patients to oscillate between supraphysiological and infraphysiological insulin doses to maintain homeostasis (10,11). Furthermore, prolonged, high-dose insulin therapy frequently predisposes patients to significant weight gain (12,13).

While establishing early and sustained glycemic control has been shown to reduce cardiovascular events in T1D patients by over 50% and prevent microvascular decay (2–4), alternative therapeutic options remain severely limited. Standard oral glucose-lowering agents like metformin fail to reduce blood glucose in T1D (10,14), and SGLT inhibitors have faced FDA approval barriers due to the heightened risk of diabetic ketoacidosis (10). Thus, discovering safer, more efficient, and insulin-independent therapies to lower glucose levels, particularly in the context of T1D, has become an urgent clinical priority.

Stearoyl-CoA desaturase-1 (SCD1) is a rate-limiting lipogenic enzyme that converts saturated fatty acids (specifically palmitate and stearate) into monounsaturated fatty acids (MUFAs, namely palmitoleate and oleate) (15,16). Previous research from our laboratory demonstrated that SCD1 deficiency not only protects against obesity and hepatic steatosis but also enhances insulin sensitivity, augments peripheral glucose uptake, and lowers blood glucose levels (15). Investigations in liver-specific SCD1 knockout (LKO) mice revealed that this increased glucose uptake occurs in both hepatic and extrahepatic tissues, mediated by a combination of insulin-dependent and insulin-independent mechanisms (15).

Additionally, we previously noted that SCD1-deficient mice exhibit elevated circulating levels of fibroblast growth factor 21 (FGF21) and adiponectin, two metabolic hormones well-known for optimizing glucose regulation (17–19). Based on these observations, we hypothesized that the alleviations in fatty liver disease and enhanced glucose clearance observed in SCD1 deficiency will prevail even in the complete absence of insulin, thereby ameliorating metabolic disease in diabetic models. In this study, we demonstrate that liver-specific SCD1 deficiency improves fatty liver disease and hyperglycemia even in the absence of insulin. The glucoregulatory benefits of SCD1 deficiency are mediated via a dual mechanism. While the upregulation of FGF21 partially contributes to improved glucose homeostasis, we show that SCD1 deficiency alters the hepatic bile acid pool, specifically elevating taurocholic acid, that sufficiently drives insulin-independent *Glut1* expression in an FXR-dependent manner. Furthermore, we show that these effects are tied to diminished oleate levels, as oleate supplementation reverses the elevated *Glut1* expression and restores hyperglycemia. Together, our results reveal that hepatic SCD1 deficiency profoundly ameliorates both hyperglycemia and fatty liver under diabetic conditions, uncovering a novel bile acid-FXR signaling axis as a promising therapeutic target for insulin-deficient states.

## Materials And Methods

### Animal studies

All experiments involving animals were conducted in accordance with the Institutional Animal Care and Use Committee guidelines of the University of Wisconsin-Madison (protocol # A005125). C57BL/6 mice were maintained in a controlled environment with a 12-hour light/dark cycle and ad libitum access to food and water. Initially, mice were fed a standard rodent chow diet (Purina 5008; Harlan Teklad, Madison, WI, USA) until they reached ∼8-10 weeks of age. For some experimental studies, mice were transferred to individual cages and fed a high-carbohydrate, very-low-fat diet (HCD; Harlan Teklad, Madison, WI, USA TD.03045) or HCD diet with triolein or tristearin for 10-14 days as indicated. The procedures of generating *SCD1^lox/lox^* (SCD1 LOX control), and *SCD1^lox/lox^*; AlbuminCre/+ (Liver specific SCD1 knockout (LKO)), SCD1 *^lox/lox^ Fgf21^loxl/lox;^ Albumin Cre/+* (DLKO) mice were previously reported (16,30). Animal experimental procedures are detailed in supplementary files.

### RNA isolation and real-time quantitative PCR

RNA was extracted from liver tissue using Trizol reagent (Life Technologies/Invitrogen, Carlsbad, CA, USA), followed by homogenization with a TissueLyzer II (Qiagen) and Turbo DNase (Ambion) treatment. cDNA synthesis was performed using a High-Capacity cDNA Reverse Transcription Kit (Applied Biosystems). Real-time quantitative PCR analysis was conducted using PowerUp SYBR Green 2x Master Mix (Thermo Fisher Scientific) on an Applied Biosystems QuantStudio 5 Real-Time PCR System in a 384-well plate. Relative gene expression was determined by the ΔΔCT method (35), with *Rps3* as a housekeeping gene. A list of primer pairs is provided in Supplementary Table S1.

RNA was extracted from HepG2 and Hepa1-6 cells using TRIzol reagent (Invitrogen), according to the manufacturer’s instructions. Maxima First Strand cDNA synthesis Kit with DNase treatment (Thermo Scientific) was used to synthesize cDNA from 1 µg of total RNA. Real-time quantitative PCR analysis was performed using PowerTrack SYBR Green master mix on a QuantStudio^TM^ 5 instrument (Thermo Scientific). Relative gene expression was determined by the ΔΔCT method (35), with *Gapdh* as a housekeeping gene. A list of primer pairs is provided in Supplementary Table S1.

### Western blotting

∼20mg of frozen liver tissue was homogenized in cold RIPA buffer (Boston Bioproducts) with protease inhibitor (Thermofisher Scientific) using a tissue lyzer (Quiagen, Hilden, Germany). Following centrifugation at 14,000 rpm at 4°C for 15 minutes, the protein amount of the resultant lysate was quantified by a BCA assay (Thermofisher Scientific). For immunoblot analysis, 20 ug of protein was solubilized in Laemmli loading buffer (Biorad) heated for 5 min at 90°C and resolved on a 4-20% acrylamide gel (Thermofisher Scientific) and subsequently transferred to a nitrocellulose membrane. Chemiluminescent detection was performed using an iBright FL1500 machine (Thermofisher Scientific), and densitometric analysis was conducted with Image J software (NIH, Bethesda, MD). Antibodies employed are listed in Supplementary Table S2.

HepG2 and Hepa1-6 cells were lysed using 200-300 µL of ice-cold Cell Lysis Buffer (Cell Signaling) with protease inhibitor cocktail (Roche) and a Bead Ruptor Elite (Omni International). Samples were centrifuged at 18,200g for 15 minutes at 4°C, and supernatants were collected. Protein concentrations were determined using BCA assay (Thermo Scientific) with BSA as a standard. Proteins were mixed with 4x NuPAGE sample buffer (Thermo Scientific) and 10x NuPAGE sample reducing agent (Thermo Scientific) and separated on precast Bis-Tris SDS-PAGE gels (Invitrogen). Proteins were transferred onto PVDF membranes using wet transfer at 30 V for 1 hour. Total protein was quantified using No-Stain protein Labeling reagent (Thermo Scientific). Membranes were blocked using 5% milk TBST for 30 minutes at room temperature before incubating with the primary antibodies diluted in 5% milk in TBST overnight at 4°C. After washing with TBST 3 times, membranes were incubated with HRP-conjugated secondary antibodies (Santa Cruz Bio Tech) diluted in 5% milk in TBST at room temperature for 1 hour. After washing with TBST 3 times, blots were developed using 1193 SuperSignal™ West Pico PLUS Chemiluminescent Substrate (Thermo Scientific). Blot images were obtained from iBright imaging system (Invitrogen). Band intensities were quantified using the iBright imaging system (Invitrogen) with automated band detection and normalized to total protein. Antibodies employed are listed in Supplementary Table S2.

### Cell culture and treatments

Hepa 1-6 and HepG2 cells were obtained from ATCC. Cells were cultured in Dulbeccos’s Modified Eagle Medium (DMEM) supplemented with 10% fetal bovine serum (FBS), penicillin (100 U/mL), and streptomycin (100 ug/mL) in a humidified atmosphere at 37°C with 5% CO_2_. One day after seeding, cells were treated with bile acid mixture mimicking either LOX or LKO hepatic bile acid pool overnight. Cells were washed with PBS once and stored in -80°C until further analysis.

### Statistical analysis

Statistical analyses were carried out in GraphPad Prism (version 10.6.1). Statistical analysis with two groups was tested by a two-tailed, Welch’s t-test, and for three or more groups was performed using one-way ANOVA. Normality tests were performed for One-way ANOVA analyses to determine whether a parametric or non-parametric test should be used. GTT analyses were subjected to Two-way ANOVA for multiple parameters. Log-rank (Mantel-Cox) test was used for survival curves.

## Results

### Hepatic SCD1 deficiency lowers systemic blood glucose levels in mice

To determine the impact of liver-specific SCD1 deficiency on systemic glucose clearance, we used liver-specific *Scd1KO* (LKO) and control (LOX) mice fed a hyperglycemia-inducing high carbohydrate, very low-fat diet (HCD) **(Fig. 1*A*)**. Oral glucose tolerance test (GTT) revealed that LKO mice exhibited lower blood glucose levels compared to LOX controls **(Fig. 1*B, C*)**. Furthermore, LKO mice presented with reduced fasting blood glucose levels and a lower HOMA IR, a surrogate marker for insulin resistance, suggesting lower insulin resistance **(Fig. 1*D, E*)**. Analysis of the glucose transporter, Glut1 (solute carrier family 2 member 1) in the liver at the transcriptional and protein level showed a significant elevation in LKO mice compared to controls, suggesting increased basal glucose intake consistent with our previous results **(Fig. 1F, G)** (15). Together, these findings demonstrate that liver-specific Scd1 deficiency induces systemic glucose clearance in the presence of a hyperglycemia-inducing diet, likely through augmented glucose uptake via Glut1 and improved insulin sensitivity.

**Fig. 1.**
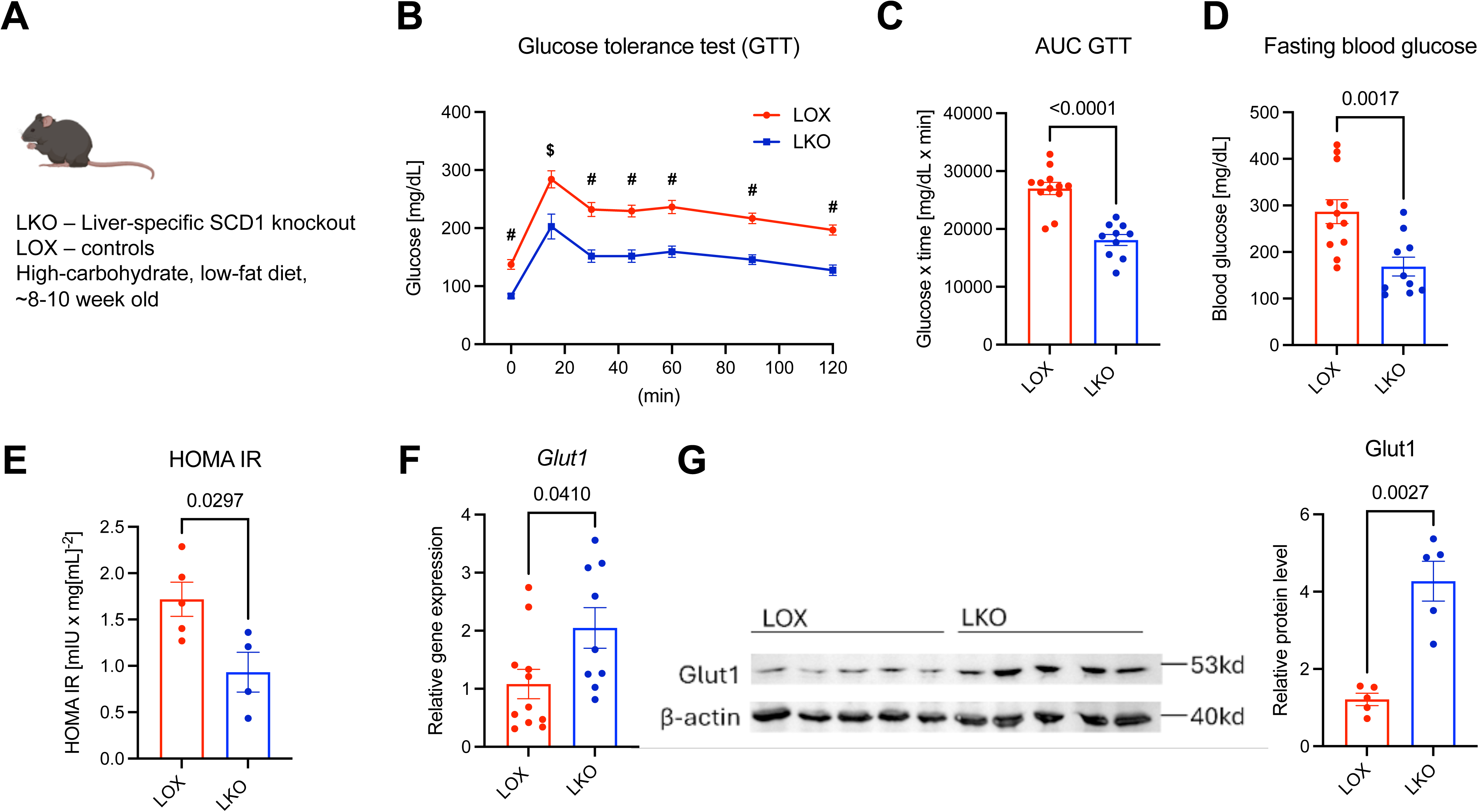
Hepatic SCD1 deficiency lowers systemic blood glucose levels in mice. A: Schematic of *in vivo* study. B: Oral glucose tolerance test (GTT) and C: Area under the curve (AUC) of the GTT in B. (LOX, *n* = 12, LKO *n* = 10. One-way ANOVA for GTT, Welch’s ttest for AUC) D: Fasting blood glucose (LOX, *n* = 12, LKO *n* = 10). E: HOMA-IR, an indicator for insulin response (LOX, *n* = 5, LKO *n* = 4). F: Relative gene expression of *Glut1* in liver (LOX, *n* = 11, LKO *n* = 9). F: Western blot of Glut1 and corresponding densitometric analysis (*n* = 5 in each group). **#***p <* 0.0001, **$***p <* 0.001. Welch’s ttest. Data are presented as mean ± SEM.

### Hepatic SCD1 deficiency reduces hyperglycemia under diabetic conditions

We have previously demonstrated that liver-specific SCD1 deficiency lowers blood glucose through increased glucose uptake via insulin-dependent and insulin-independent mechanisms (15). To further delineate the role of insulin-independent mechanism in glucose regulation, we employed a streptozotocin-induced diabetes model. Briefly, LOX and LKO mice were fed HCD for a week, followed by an intraperitoneal (IP) injection with streptozotocin (20). Streptozotocin is selectively imported in pancreatic beta cells via Glut2 and induces diabetes by causing dysfunction of the pancreatic beta cells (21). Consistently, streptozotocin-treated LOX control mice exhibited overt diabetes characterized by pronounced hyperglycemia only two days post-streptozotocin administration, with blood glucose levels surpassing twice the prediabetic threshold **(Fig. 2*A*)**.

**Fig. 2.**
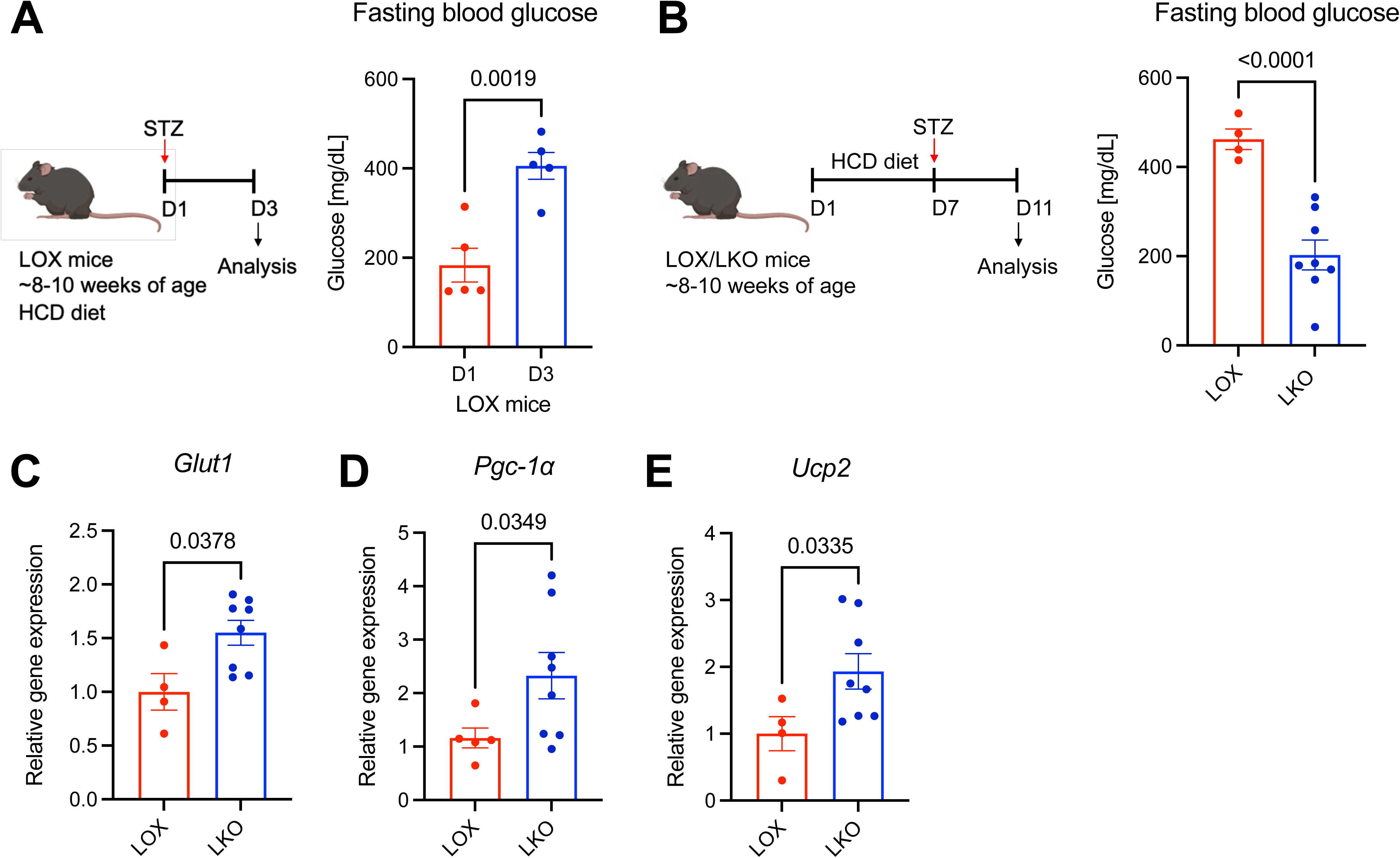
Hepatic SCD1 deficiency lowers blood glucose levels independent of insulin in diabetic mice. A: Schematic of *in vivo* study and fasting blood glucose of LOX mice treated with streptozotocin (STZ) for 2 days. *n* = 5 mice per group. Welch’s ttest. B: Schematic of *in vivo* study and fasting blood glucose levels of STZ-treated diabetic mice. C-E: Liver gene expression of B: *Glut1,* C: *Pgc1α,* and D: *Ucp2*. *n* = 4-8 mice per group. Welch’s ttest. Data are presented as mean ± SEM.

Remarkably, mice with liver-specific Scd1 deficiency were protected against streptozotocin-induced hyperglycemia. Our results revealed that LKO mice had significantly lower fasting blood glucose levels, approximately 50% that of LOX levels on day 5 post-streptozotocin administration **(Fig. 2*B*)**. We also observed an increase in liver *Glut1* gene expression **(Fig. 2*C*)**. This suggests that even under diabetic conditions, LKO mice induce systemic glucose clearance, potentially via Glut1. Additionally, we observed enhanced expression of *Pgc1a* and *Ucp2* in the liver, which is associated with thermogenesis **(Fig. 2*D, E*)**, consistent with previous findings in non-diabetic LKO mice (16). Together, these findings indicate that hepatic SCD1 deficiency ameliorates hyperglycemia under diabetic conditions independent of insulin.

### Hepatic SCD1 deficiency protects against fatty liver under diabetic conditions

We have previously found that liver-specific Scd1 deficiency protects mice against fatty liver when fed a HCD through the downregulation of *de novo* lipogenesis (16). In this study, we tested whether this protective effect persists under diabetic conditions. Histological analysis of liver tissues revealed a striking decrease in lipid droplets in LKO mice compared to LOX mice in diabetic conditions (**Fig. 3*A*, *B*).** Analysis of gene expression changes related to *de novo* lipogenesis and lipid droplet formation corroborated these results. Transcript levels of perilipin 2 (*Plin2*), acetyl-CoA carboxylase (*Acc*), elongation of very long chain fatty acids-like 6 (*Elovl6*), sterol regulatory element-binding protein-1c (*Srebp1c*), and fatty acid synthase (*Fas*) were all downregulated in LKO mouse livers compared to LOX mice (**Fig. 3*C-G*)**. Overall, these findings indicate that Scd1 deficiency in the liver protects mice against fatty liver disease independent of insulin.

**Fig. 3.**
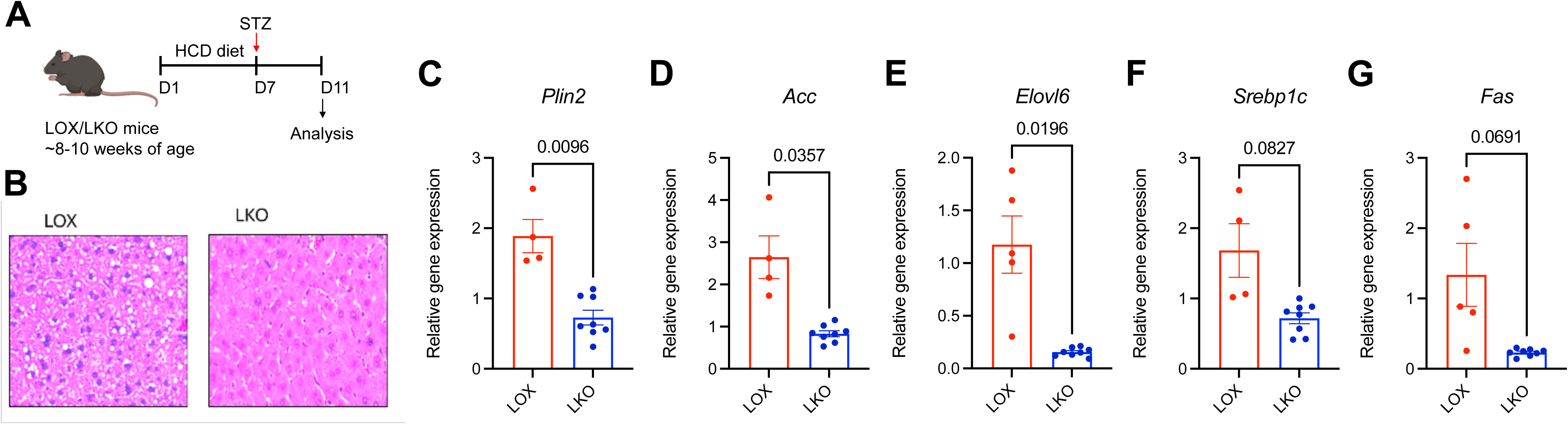
Hepatic SCD1 deficiency prevents fatty liver development under diabetic conditions. A: Schematic of the *in vivo* study. B: H&E staining of the liver in LOX and LKO mice. C-G: Relative expression of genes related to lipogenesis and lipid droplet formation. *n* = 4-8 mice per group. Welch’s ttest. Data are presented as mean ± SEM.

### Hepatic SCD1 deficiency leads to an increase in liver and plasma bile acids

We next sought to uncover how depletion of hepatic Scd1 leads to an increase in Glut1 expression in the liver and amelioration of systemic hyperglycemia. Whole-body *Scd1KO* results in reduced cholesterol-derived lipid synthesis, leading to higher free cholesterol levels (22). We hypothesized that a liver-specific *Scd1KO* would also result in elevated free cholesterol, which would be diverted toward hepatic cholesterol-derived bile acid (BA) synthesis (23). Consistent with our hypothesis, we found a significant 2-fold elevation in total liver BAs, and a striking 10-fold elevation in total plasma BAs in LKO mice compared to LOX (**Fig. 4*A-C***). Hepatic and plasma levels of both unconjugated and conjugated primary bile acids were elevated in LKO mice. Secondary BAs were absent in the liver, and present in low levels in plasma, with a significant increase in deoxycholic acid (DCA) levels in LKO mice, but no change in levels of ursodeoxycholic acid (UDCA) and lithocholic acid (LCA) in LKO mice compared to LOX. Taurocholic acid (TCA) emerged as the most predominant and significantly elevated BA species in LKO liver and plasma compared to LOX mice.

**Fig. 4.**
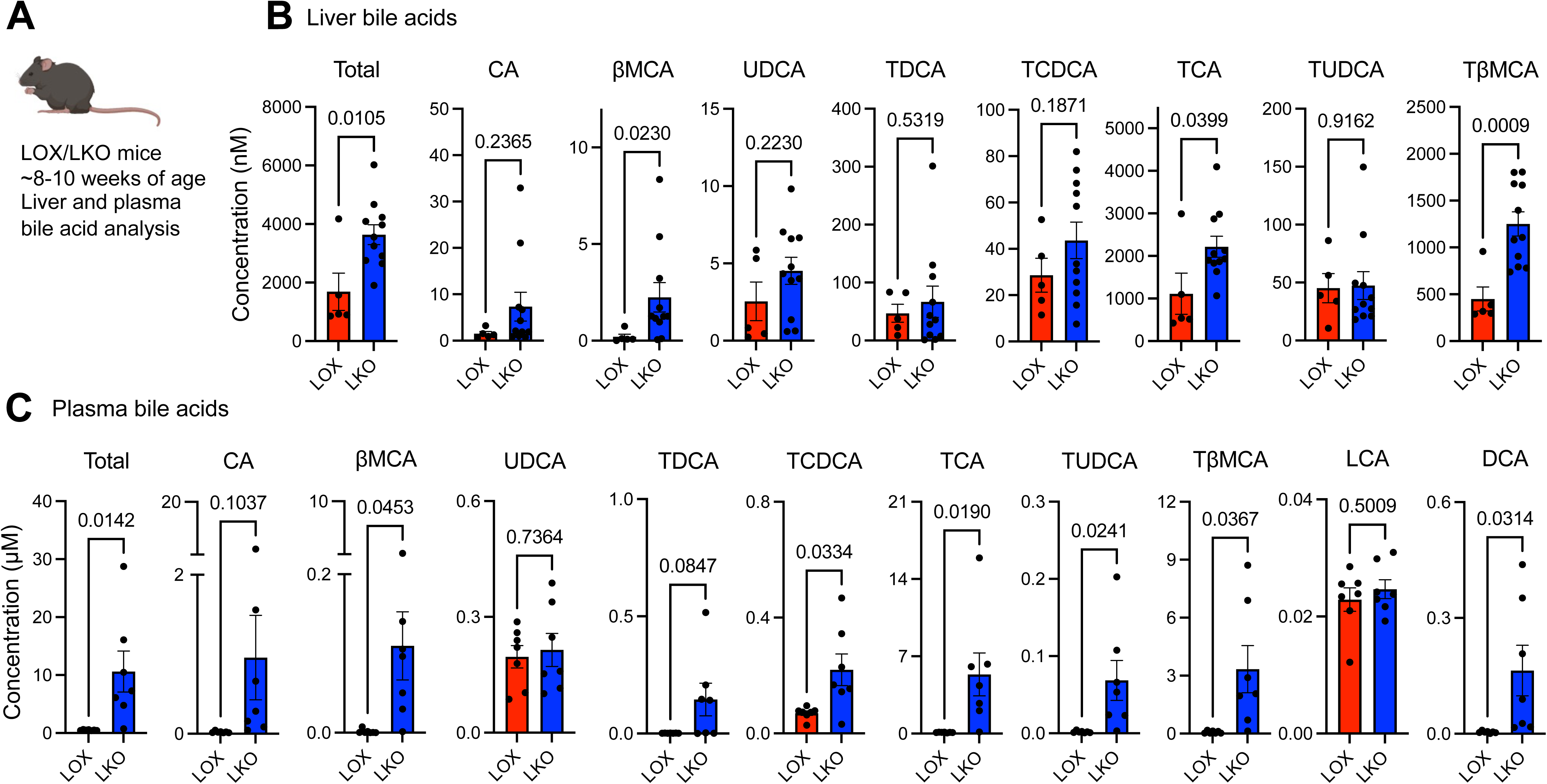
Liver and plasma BAs are elevated in LKO mice. A: Schematic of *in vivo* study. B: Liver BAs. LOX, *n* = 5; LKO *n* = 11 mice. C: Plasma BAs. *n* = 7 mice in each group. Welch’s ttest. Data are presented as mean ± SEM. CA – cholic acid; βMCA – beta muricholic acid; UDCA – ursodeoxycholic acid; TDCA – tauro-deoxycholic acid; TCDCA – tauro-chenodeoxycholic acid; TCA – tauro-cholic acid; TUDCA – tauro-ursodeoxycholic acid; TβMCA – tauro-beta muricholic acid; LCA – lithocholic acid; DCA – deoxycholic acid.

### Hepatic BAs shifted in LKO mice increase *Glut1* expression in the liver via FXR

In recent years, BAs have gained significant attention as modulators of metabolism and disease, specifically diabetes (24,25). We hypothesized that BAs shifted in LKO mice trigger an increase in *Glut1* expression in the liver. To test this hypothesis, we generated an *in vitro* pool of BAs that match the average physiological concentrations observed in LKO and LOX mouse liver samples **(Fig. 4*B*, 5*A*)**^27-29^. We treated human HepG2 liver cells and mouse Hepa1-6 liver cells with these pools of BAs to measure *Glut1* expression. Strikingly, LKO pool of BAs increased *Glut1* expression compared to the LOX pool **(Fig. 5*B, C*)**.

**Fig. 5.**
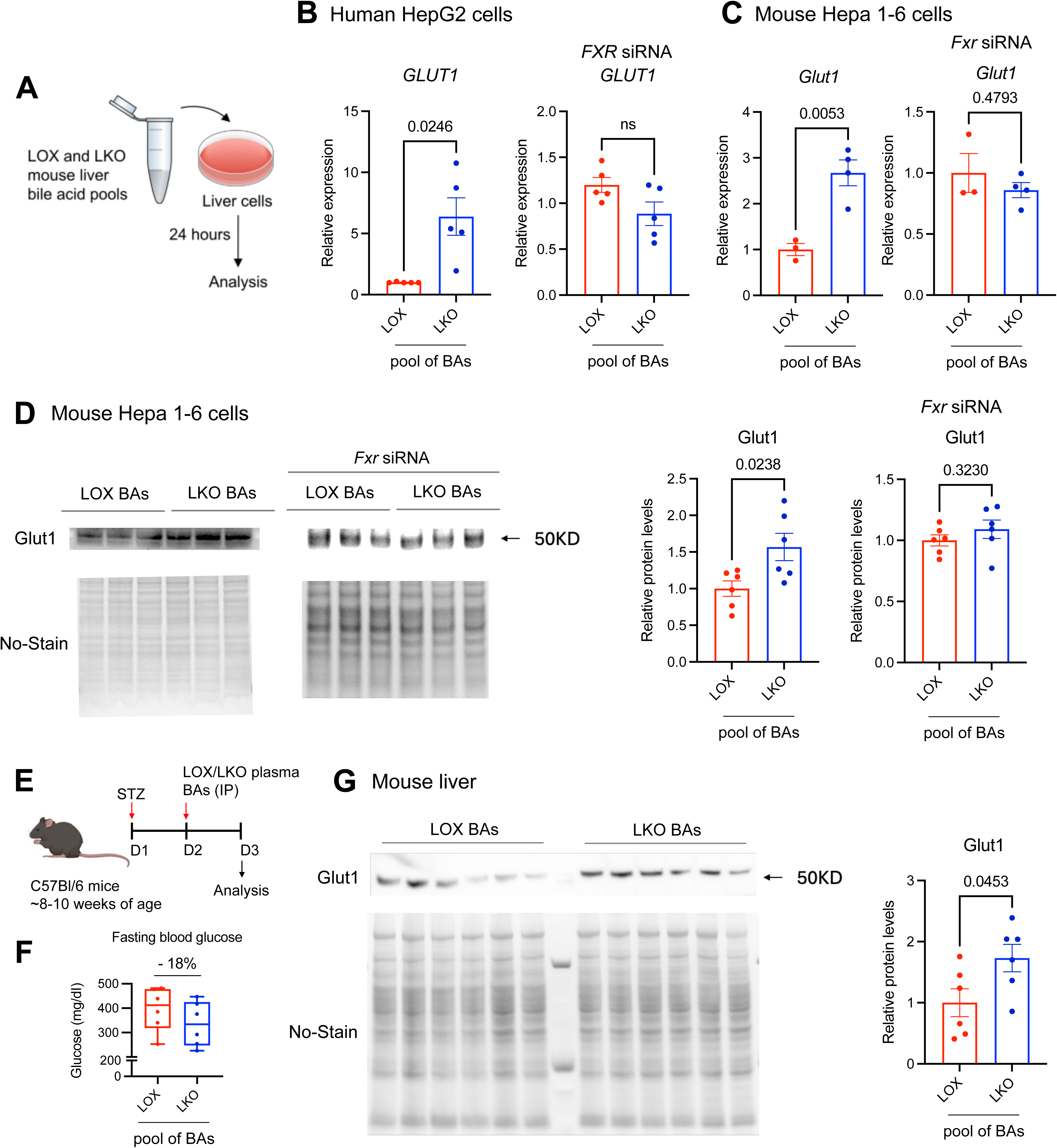
LKO liver BAs increase *Glut1* expression in the liver via FXR. A: Schematic of *in vitro* experiment. B, C: *GLUT1* expression in B: HepG2 and C: Hepa 1-6 cells treated with LOX and LKO liver BA pools with or without *FXR* siRNA. B, *n* = 5 in each group. C: *n* = 3 in LOX, *n* = 4 in LKO group. D: Western blot analysis of Glut1 in mouse Hepa1-6 cells with or without *Fxr* siRNA. *n* = 6 in each group. E: Schematic of *in vivo* experiment. F: LKO plasma BA treated mice show a 18% reduction in fasting blood glucose. G: Western blot analysis of Glut1 in mouse liver samples. *n* = 6 in each group. Welch’s ttest. Data are presented as mean ± SEM.

Bile acids modulate gene transcription primarily by activating nuclear hormone receptors, most notably the farnesoid X receptor (FXR) (26). FXR agonism has been shown to reduce hyperglycemia and fatty liver disease in animal models (27). TCA is a strong FXR agonist, known to improve metabolic disease via FXR agonism in many organs (28). Considering that TCA was the most predominant BA in the liver and plasma, and was significantly elevated in LKO mice, we hypothesized that Fxr agonism in liver cells increases *Glut1* expression. To test this hypothesis, we performed an siRNA-mediated knockdown of *FXR* in human HepG2 and mouse Hepa 1-6 liver cells, followed by administration of LOX and LKO liver BA pools. Strikingly, LKO BA-mediated increase in GLUT1 was abolished in the *FXR* siRNA-treated liver cells, demonstrating that FXR is required for LKO hepatic BAs to induce *GLUT1* expression **(Fig. 5*B, C*)**. Western blot analysis of Glut1 in mouse Hepa 1-6 cells further corroborated our qPCR results, overall suggesting that BAs shifted in the liver due to Scd1 deficiency can induce *Glut1* expression via agonism of Fxr **(Fig. 5*D*)**.

### Plasma BAs shifted in LKO mice increase hepatic Glut1 expression under diabetic conditions

To test whether BAs shifted in LKO mice can induce *Glut1* expression systemically, we administered LOX and LKO plasma BAs pools to diabetic mice. Briefly, wild-type mice were treated with streptozotocin to deplete pancreatic beta cells, followed by administration of LKO and LOX plasma BA pools via IP injection **(Fig. 5*E*)**. LKO plasma BA treated animals showed a 18% decrease in fasting blood glucose levels within 1 day of LKO BA treatment compared to LOX BA pools **(Fig. 5*F*)**. Strikingly, hepatic Glut1 protein levels were significantly elevated in the LKO BA treatment, suggesting that BAs shifted in Scd1-deficient animals can induce *Glut1* expression in the liver in an insulin-independent manner.

### Improvement in glucose uptake under SCD1 deficiency is partially mediated by FGF21

Our studies have shown that hepatic Scd1 ablation ameliorates hyperglycemia both in healthy and diabetic mice subjected to a hyperglycemia inducing diet. Intriguingly, our previous studies uncovered a substantial increase in FGF21, an FXR-induced hepatokine that improves glucose metabolism in the livers of LKO mice (15,29). To probe the role of FGF21 in glucose metabolism in LKO mice, we used mice with a double knockout of Scd1 and FGF21 in the liver (DLKO) (30) administered a HCD **(Fig. 6*A*)**. DLKO mice showed a significant increase in fasting blood glucose levels compared to LKO mice, reaching levels comparable to LOX mice, suggesting that loss of FGF21 abolishes the glucoregulatory benefits of hepatic Scd1 deficiency **(Fig. 6*B*)**. Loss of FGF21 also diminished insulin response in DLKO mice **(Fig. 6*C*)**. However, an oral glucose tolerance test revealed that while LKO mice were efficient in systemic glucose clearance, DLKO mice showed only a partial defect in reducing hyperglycemia as compared to LOX mice **(Fig. 6*D, E*)**. This suggests that FGF21 partially contributes to improved glucose homeostasis in LKO mice. Consistently, DLKO mice had a partial increase in *Glut1* expression compared to LOX mice **(Fig. 6*F*)**. These findings indicate that hepatic FGF21 plays a significant, yet partial role in modulating glucose levels under hepatic Scd1 deficiency, suggesting that other factors downstream of Fxr may be involved in LKO glucoregulatory effects either independently or synergistically with FGF21.

**Fig. 6.**
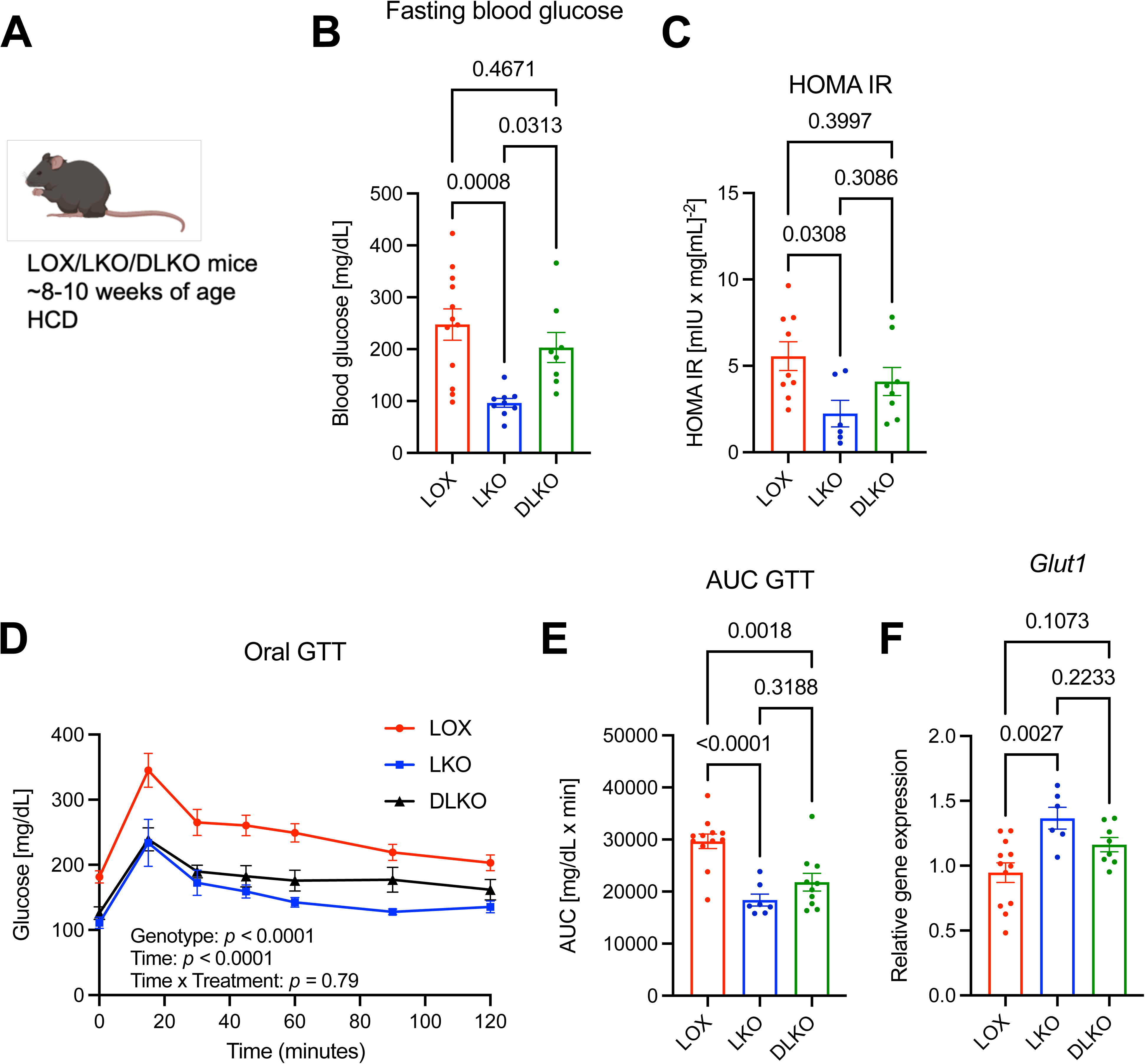
Hepatic SCD1 deficiency lowers systemic blood glucose levels partially through FGF21. A: Schematic of *in vivo* study. B: Fasting blood glucose (LOX, *n* = 12, LKO *n* = 9, DLKO *n* = 8). C: HOMA-IR, an indicator for insulin response (LOX, *n* = 9, LKO *n* = 6, DLKO *n* = 8). D: Oral glucose tolerance test (GTT) and E: Area under the curve (AUC) of the GTT in D. (LOX, *n* = 12, LKO *n* = 6, DLKO *n* = 8. One-way ANOVA for GTT, Welch’s ttest for AUC). F: Relative gene expression of *Glut1* in liver (LOX, *n* = 11, LKO *n* = 9). F: Glut1 expression (LOX, *n* = 12, LKO *n* = 6, DLKO *n* = 8). One way ANOVA. Data are presented as mean ± SEM.

### Oleate restores hyperglycemia in liver-specific Scd1-deficient mice under diabetic conditions

Scd1 catalyzes the delta-9 desaturation of fatty acids, mainly palmitate and stearate to palmitoleate and oleate, respectively **(Fig. 7*A*)** (16,31–33). We sought to determine whether oleate, a product of Scd1, or stearate, the substrate of Scd1 can modulate blood glucose levels under diabetic conditions. To test this, we fed streptozotocin-induced diabetic LOX and LKO mice fed a HCD diet supplemented with either stearate (in the form of tristearin) or oleate (in the form of triolein) **(Fig. 7*B*)**. Stearate drastically lowered the survival rate of the LOX control mice to a mere 25% by the end of the experiment **(Fig. 7*C*)**, prompting their exclusion from subsequent analyses due to the limited number of surviving animals. In contrast, LKO mice showed protection against stearate toxicity with a survival of 75% **(Fig. 7*C*)**. Supplementation with oleate allowed the survival of both LOX and LKO mice **(Fig. 7*C*)**.

**Fig. 7.**
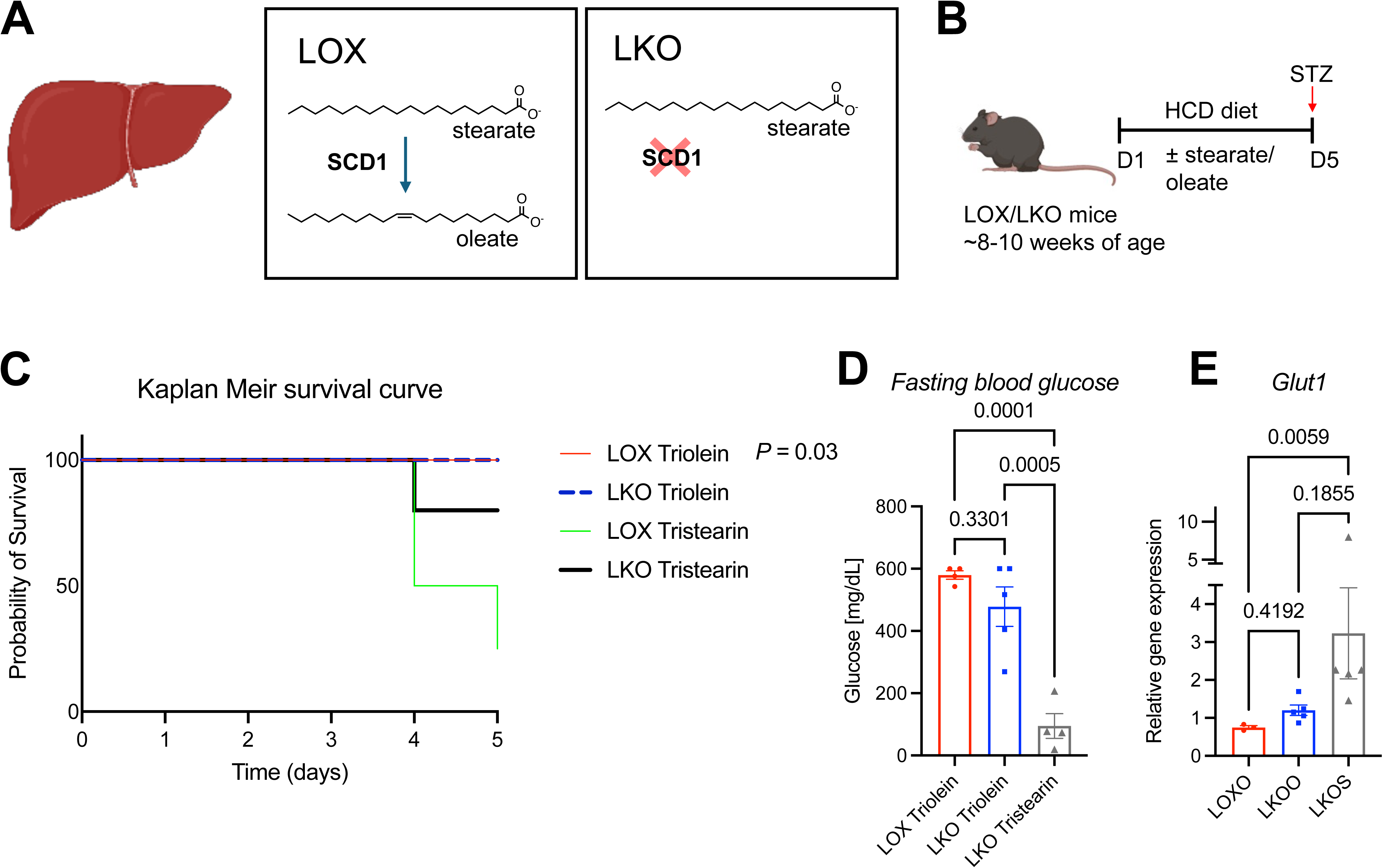
Oleate restores hyperglycemia in liver-specific Scd1 deficient diabetic mice. A: Schematic of Scd1 enzyme action in liver. B: Schematic of *in vivo* experiment. C Kaplan Meir survival curve. (LOX Triolein *n* = 4, LKO Triolein *n* = 5, LOX Tristearin *n* = 4, LKO Tristearin *n* = 5. Log-rank (Mantel-Cox) test). D: Fasting blood levels. (LOX Triolein *n* = 4, LKO Triolein *n* = 5, LKO Tristearin *n* = 5. One-way ANOVA). E: Relative liver gene expression of *Glut1* (LOX Triolein *n* = 4, LKO Triolein *n* = 5, LKO Tristearin *n* = 5. One-way ANOVA). Data are presented as mean ± SEM.

Remarkably, oleate supplementation in LKO mice restored the blood glucose levels to those observed in LOX mice treated with oleate **(Fig. 7*D*)**. In contrast, stearate supplementation maintained low blood glucose levels in Scd1 deficient mice **(Fig. 7*D*)**, indicating a distinct metabolic response to the fatty acids. Moreover, the supplementation of oleate but not stearate attenuated the elevated hepatic *Glut1* gene expression observed in LKO mice **(Fig. 7*E*)**. These findings indicate that diminished oleate levels play a role in lowering blood glucose under hepatic Scd1 deficiency, and replenishing oleate restores the blood glucose, potentially influencing glucose uptake. This oleate-specific effect underscores the unique metabolic consequences of fatty acids modulated in the liver by Scd1.

## Discussion

The critical challenge in managing Type 1 Diabetes (T1D) remains an absolute reliance on exogenous insulin therapy. This monotherapy is clinically restricted by a narrow therapeutic window, high glycemic variability, and secondary complications such as severe hypoglycemia and weight gain. Identifying physiological pathways capable of stimulating blood glucose clearance entirely independent of insulin action is therefore of paramount clinical importance. In this study, we demonstrate that liver-specific stearoyl-CoA desaturase-1 (SCD1) deficiency profoundly protects against diabetic hyperglycemia and hepatic steatosis through an insulin-independent mechanism coordinated by endogenous bile acid (BA) signaling. While our previous work established that SCD1 deletion increases systemic glucose disposal via both insulin-dependent and insulin-independent mechanisms (15), our present findings reveal that the insulin-independent branch is driven by BAs activating farnesoid X receptor (FXR) and upregulating glucose transporter 1 (*Glut1*) expression.

Under physiological conditions, the liver primarily acts as a glucose-exporting organ during fasting, while postprandial hepatic glucose uptake relies heavily on insulin-dependent pathways. However, patients receiving subcutaneous insulin therapy suffer from a lack of portal-hepatic insulin normalization; the liver receives insulin at substantially lower concentrations than are physiologically required to suppress hepatic glucose output and optimize clearance. Inhibiting hepatic SCD1 bypasses this clinical barrier by upregulating *Glut1*, a glucose transporter that facilitates baseline, non-insulin-dependent glucose uptake. Mechanistically, we show that this upregulation is driven by alterations in the hepatic lipidome. Whole-body SCD1 deficiency is known to impair cholesterol-derived lipid synthesis, accumulating free cholesterol. (34) Our data demonstrates that liver-specific SCD1 deficiency redirects this excess cholesterol into alternative metabolic pathways, triggering a significant 2-fold elevation in total liver BAs and a striking 10-fold increase in plasma BAs.

Among these species, taurocholic acid (TCA), a potent endogenous agonist of the nuclear hormone receptor FXR, emerged as the predominant elevated species. Our *in vitro* siRNA knockdown assays confirmed that this specific LKO bile acid pool directly induces *Glut1* expression in an FXR-dependent manner. Furthermore, the immediate 18% drop in fasting blood glucose observed after administering this LKO plasma BA pool to streptozotocin-induced diabetic mice provides *in vivo* validation that shifting endogenous bile acid composition is sufficient to execute insulin-independent systemic glucose clearance.

Our previous working hypothesis focused on the PGC-1α–FGF21 and adiponectin endocrine axes as primary drivers of glucose management in SCD1-deficient models. Deletion of hepatic *Fgf21* in LKO mice certainly aggravated fasting glucose levels and partially weakened insulin sensitivity. However, during oral glucose challenge, DLKO mice retained a robust, persistent capacity for systemic glucose clearance, showing only a partial defect compared to controls. Similarly, *Glut1* expression was only partially blunted in these models. These findings indicate that while FGF21 (which is itself an FXR-induced hepatokine) acts as a cooperative systemic metabolic optimizer, the local hepatic bile acid shift and autonomous FXR activation are powerful enough to independently sustain basal glucose transport. This underscores a redundant biological framework designed to defend glycemic homeostasis under profound metabolic stress.

One of the most striking findings of this study is that dietary supplementation with oleate, the direct monounsaturated product of SCD1 desaturation, completely reversed the therapeutic effects of SCD1 deficiency, suppressing hepatic *Glut1* expression and restoring diabetic hyperglycemia. Conversely, supplementation with the substrate stearate left the low-glycemic, high-*Glut1* phenotype entirely intact. This reveals that the upstream metabolic sensor governing cholesterol redirection and bile acid synthesis is remarkably sensitive to the cellular monounsaturated-to-saturated fatty acid ratio. Furthermore, we observed a dramatic protective effect against saturated fat lipotoxicity. While excess stearate administration proved lethal to 75% of control mice, LKO mice were largely protected from this toxicity. Coupled with the elevated hepatic expression of *Pgc1α* and *Ucp2*, these data suggest that SCD1 deficiency actively forces the liver to redirect potentially hazardous, lipotoxic saturated substrates away from harmful *de novo* lipogenesis, which we show is thoroughly suppressed, and into protective thermogenic dissipation and bile acid synthesis pathways. This multi-pronged metabolic rewiring simultaneously neutralizes diabetic fatty liver disease and clears circulating glucose.

By targeting a single lipogenic node, hepatic SCD1 ablation introduces a therapeutic strategy capable of concurrently mitigating hyperglycemia and hepatic steatosis - two of the most debilitating and interlinked comorbidities in diabetes care. For T1D patients, capitalizing on an endogenous bile acid-FXR signaling axis to optimize hepatic basal glucose uptake could drastically lower exogenous insulin requirements. This could alleviate the day-to-day burden of frequent injections and significantly minimize long-term risks of insulin-induced weight gain and macrovascular decline.

While our human HepG2 cell assays demonstrate that human liver lines share this FXR-dependent *Glut1* response to the LKO bile acid profile, translating these findings to human therapeutics requires careful consideration. Mouse and human bile acid pools possess inherent compositional disparities (e.g., the prevalence of hydrophilically distinct muricholic acids in rodents). Future investigations should prioritize validating these specific FXR-binding kinetics utilizing human-prevalent bile acid profiles and evaluating whether circulating plasma BAs similarly activate peripheral or extrahepatic glucose disposal pathways.

Nonetheless, our findings fundamentally establish hepatic SCD1 and its downstream bile acid-FXR signaling networks as a compelling, insulin-independent therapeutic framework for the management of insulin-deficient diabetes.

## Acknowledgments

This study was supported by the National Institutes of Health (R01 DK118093 to J.M.N., R00 DK128503 to S.N.C., T32 GM152341 to W.S.K., T32 AG000213 to N.W., and T32 GM135066 to J.W.), the Wisconsin Alumni Research Foundation, and the Department of Biochemistry at UW-Madison.

## Article Information

### Author Contributions

Conceptualization and study design was carried out by M.K., S.N.C., and J.M.N. All *in vivo* experiments were performed by M.K., D.K., W.S.K., V.P.C., E.A., J.R.M., D.C., K.G., and D.B., including animal breeding, maintenance, experiments, data collection, and end-point analyses. N.W., and J.W. developed and optimized bile acid LC-MS analysis. M.K., and D.K. performed bile acid LC-MS analyses from *in vivo* samples, and all *in vitro* experiments. M.K. and S.N.C. wrote the paper and prepared all figures. All authors edited and contributed to the critical review of the paper.

### Data and Resource Availability

All data pertaining to the study are reported in the manuscript and associates supplemental files. Additional data or information is available upon request. Dr. Snehal N. Chaudhari and Dr. James Ntambi are the guarantors of this work and, as such, had full access to all the data in the study and takes responsibility for the integrity of the data and the accuracy of the data analysis.

