## Supplementary tables for "Hepatic stearoyl-CoA desaturase deficiency ameliorates hyperglycemia through bile acid signaling in an insulin-independent manner"

and

James M. Ntambi PhD
University of Wisconsin-Madison
433 Babcock Drive, Madison, WI 53706


**Table S1. Primers**

| Name (m=mouse; h=human) | Forward (5' – 3') | Reverse (5' – 3') |
|---|---|---|
| m-Glut1 | TATCAGCCACTCTCCTATCTCC | AGGTCCAGCCCTACAGATTA |
| m-Pgc1-a | TGTCGCCTTCTTGCTCTTCC | GGAACACGACCTGTGTCGAG |
| m-Ucp2 | CCAGCCTACAGATGTGGTAAA G | TCGACAGTGCTCTGGTATCT |
| m-Srebp1c | CCATTGACAAGGCCATGCAG | TTGCTGGTACCGTGAGCTAC |
| m-Elovl6 | GAACAAGCGAGCCAAGTTTG | TGTAAGCACCAGTTCGAAGAG |
| m-Fas | GCTGCGGAAACTTCAGGA AAT | AGAGACGTGTCACTCCTGGACTT |
| m-Acc | TGACAGACTGATCGCAGAGAAAG | TGGAGAGCCCCACACACA |
| m-Plin2 | CGTCTGTCTGGACCGAATAAA | CACACGCCTTGAGAGAAACA |
| m-Fgf21 | CTACACAGATGACGACCAAGA | CTTTGAGCTCCAGGAGACTTT |
| m-Fxr | AATCTCTTCCCAGCAGCCTTTG | CATCCGAACTTTAGCCAGCC |
| m-18S | ATTGGAGCTGGAATTACCGC | CGGCTACCACATCCAAGGAA |
| h-Glut1 | TGGCATCAACGCTGTCTTCT | CTAGCGCGATGGTCATGAGT |
| h-Fxr | ACTGACCTGTGAGGGGTGTA | ACATACATTCAGCCAACATTCC |
| h-Gapdh | F-GAAGGTGAAGGTCGGAGT | CATGGGTGGAATCATATTGGAA |

**Table S2. Antibodies**

| **Name** | **Citation** | **Supplier** | **Cat no.** |
|---|---|---|---|
| GLUT1 [EPR3915] | PMID: 35263587 | Abcam | **ab115730** |
| GLUT1 | | Proteintech | **21829-1ap** |
| β-actin | PMID: 37549300 | Thermofisher | **PA1183** |
| HRP-goat anti-rabbit IgG | PMID: 37783697 | Cell | **7074** |
